# Riverine Invertebrates Exhibit Spatial Asymmetry in Temporal Ecological Processes Across Landscapes

**DOI:** 10.1101/2024.08.30.610482

**Authors:** Xiaowei Lin, Jialu Wan, Qingyi Luo, Qinghua Cai, Ming-Chih Chiu, Vincent H. Resh

**Author notes:** Corresponding Author: **MCC:**. **Conflict of Interest:** The authors declare no conflict of interest. **Author Contributions:** XL, QL, QC, MCC, and VHR drafted the article and conceptualized the project. XL and MCC devised the research methodology. XL, JW, and MCC conducted data processing and analysis. All authors reviewed and revised the article. **Data availability statement:** The data that supports the findings of this study are available in the supporting information Data S4 at https://onlinelibrary.wiley.com/doi/10.1111/jbi.13913. **Funding:** Qinghua Cai was funded by the Second Tibetan Plateau Scientific Expedition and Research Program (2019QZKK0402). Ming-Chih Chiu was funded by the Chinese Academy of Sciences Taiwan Young Talent Programme (Grant No. 2017TW2SA0004-Y-Y).

## Abstract

The environment and its variability exhibit spatial heterogeneity in influencing ecological processes and community dynamics across landscapes. However, the mechanisms remain unexplored.

This study investigates relative significance of temporal ecological processes that shape invertebrate community dynamics across 15 rivers in the European Iberian Peninsula over 21 years.

Spatial asymmetry in community dynamics was driven by a combination of temporal deterministic and stochastic processes (i.e., environmental filtering and temporal distance within the same locations, respectively).

It is noteworthy that the relative importance of deterministic versus stochastic processes diminishes with increasing elevation.

The analysis of community dynamics in diverse landscapes offers a foundation for anticipating and mitigating the consequences of prospective environmental transformation on biodiversity, thereby directing efficacious conservation strategies.

## 1 Introduction

Overwhelming evidence suggests an unprecedented decline in global biodiversity and the vital ecosystem services it delivers (Liu et al. 2023). This trend is particularly pronounced for species facing varied fates due to adaptations in their life histories, morphology, and physiology amidst the complexities of a changing global environment. For instance, species inhabiting high-altitude or -latitude regions often struggle to adapt to shifting environmental conditions (Doxa et al. 2022). Spatial heterogeneity within landscapes is crucial for understanding how diverse biological communities respond to both anthropogenic and natural environmental changes through various ecological processes.

Here, we propose that spatial asymmetry of the environmental variability play a key role in shaping the geographic distribution of temporal ecological processes (i.e., within the same locations) that govern community dynamics. Specifically, the influence of the environmental variability on community dynamics has been extensively documented (Hiltunen et al. 2015, Zhang et al. 2022, Lin et al. 2024a). Additionally, with environmental variability intensifying at higher latitudes and elevations (Antunez 2023, Lin et al. 2024a), disparities in community dynamics may arise due to geographical distinctions. Existing research has predominantly focused on two main areas: the spatial patterns of environmental variability (Kozlovsky et al. 2018, Whitenack et al. 2023) and the impacts of these variability on biogeographic patterns and community dynamics (Chan et al. 2018, Bier et al. 2022, Wayman et al. 2022). However, the interaction between spatial patterns of environmental variability and the geographic distribution of community dynamics remains largely unexplored from a mechanistic perspective involving temporal ecological processes. This current study has a corresponding bioRxiv preprint (Lin et al. 2024b), which has been credited and expanded upon by our another work that examines whether changing climate alters these spatial patterns of temporal ecological processes (Lin et al. 2024a).

Community dynamics studies provide different insights compared to those that examine spatial interactions between communities. Therefore, community dynamics research is gaining importance (e.g. Magurran et al. 2019, Wu et al. 2022, Lin et al. 2024a). These studies explore how both temporal deterministic processes and stochastic processes (e.g., ecological drift) influence community dynamics (Dini-Andreote et al. 2015, Gao and Liu 2018, He et al. 2021). Additionally, the importance of temporal ecological processes is integral to understanding the underlying mechanisms that result in the resulting spatial patterns in community dynamics (Dini-Andreote et al. 2015, Bontemps et al. 2024). For instance, the temporal deterministic processes tend to prevail over longer time scales, influencing community dynamics (Vanwonterghem et al. 2014, Aguilar and Sommaruga 2020).

To fill the aforementioned gap, this study represents an effort to investigate the intricate relationship between the spatial aspects of both environmental variability and community dynamics from a mechanistic standpoint rooted in temporal ecological processes. We analyzed the ecological processes influencing community dynamics using extensive long-term data on riverine invertebrates from a geographically expansive and continuously monitored area. The present study postulates that spatial patterns of environmental variability can exert a considerable influence on the geographic patterns observed in community dynamics. Furthermore, we assessed how elevation changes might modify the relative contributions of stochastic and deterministic processes driving biological community dynamics.

## 2 Materials and Methods

### 2.1 Study area and macroinvertebrate data

The benthic macroinvertebrate data used in this study were originally collected by Cañedo-Argüelles et al. (2020) from 15 rivers located in the northeastern part of the Iberian Peninsula in Spain (Fig. 1). This region is characterized by a Mediterranean climate, with annual precipitation ranging from 500 to 1400 mm and mean annual temperatures between 7 and 15°C. According to the Assessment Criteria of Mediterranean Rivers (Sanchez-Montoya et al. 2009), the human pressures in the studied river reaches are assessed as relatively low.

**Figure 1.**
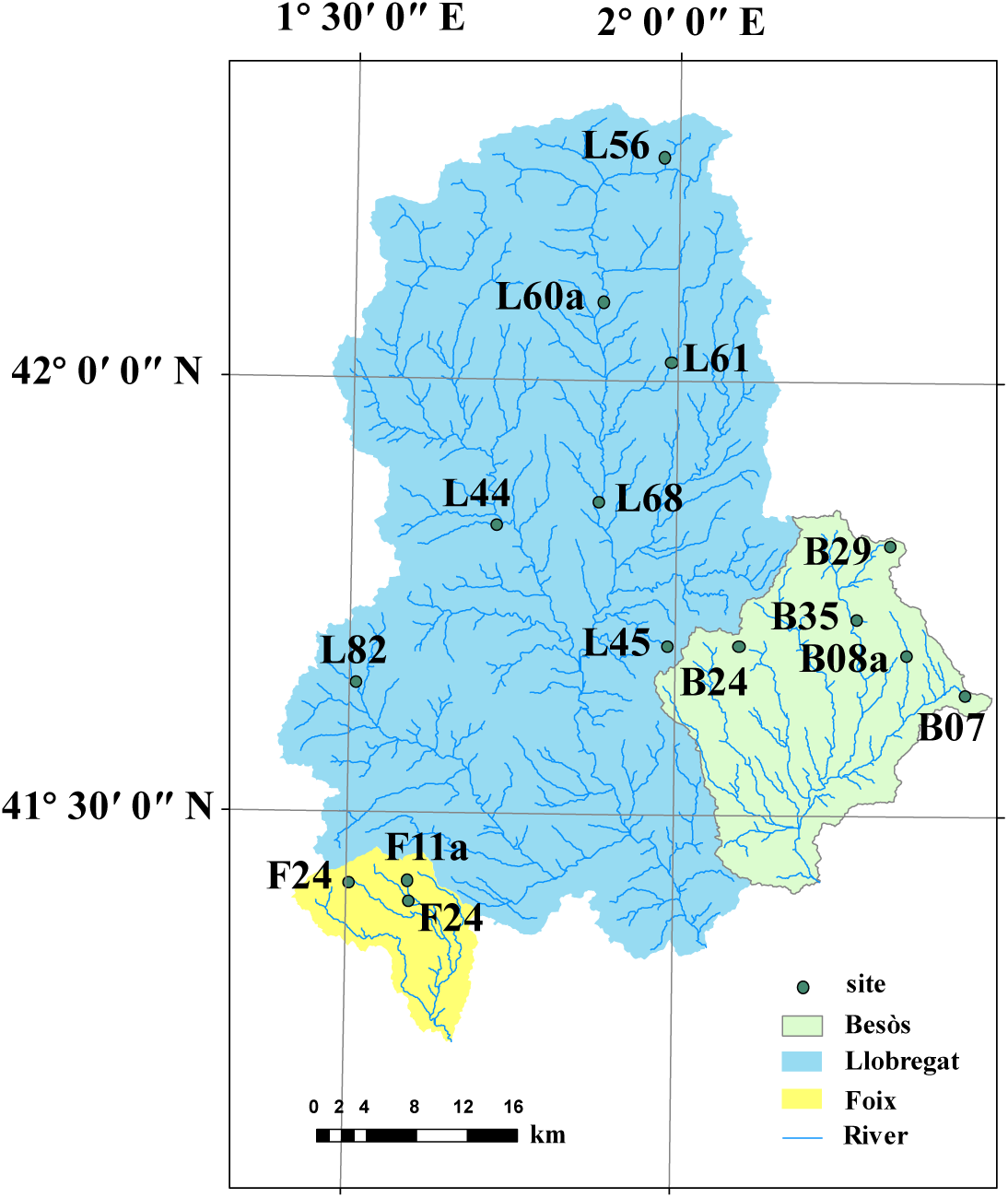
Sampling sites in the Iberian Peninsula in Spain.

### 2.2 Environmental variables and time variables

We use water quality and climate variables to represent the temporal deterministic processes. In the absence of complete data on internal variables pertaining to the stream, such as water quality, we employed the Multiple Imputation Through XGBoost (mixgb) method to address the issue of missing data. The method employs XGBoost’s gradient boosting tree algorithm, which enables the model to discern intricate non-linear relationships and interactions between variables, thereby enhancing accuracy. Moreover, it employs a combination of subsampling and predictive mean matching to minimise bias and facilitate the capture of variation. Please refer to Appendix S3 for details. The aforementioned operations are conducted with the ‘mixgb’ package in R (Deng 2023).

Monthly climatic variables such as mean minimum temperature, mean maximum temperature, and total precipitation were sourced from the WorldClim 2 database (Fick and Hijmans 2017) for each sampling site over the period spanning 1997 to 2017. These data were further processed using the “biovars” function within the R package “dismo” (Hijmans 2023) to derive 19 physical and climatic variables, including annual precipitation, mean annual temperature, minimum temperature of the coldest month, and maximum temperature of the hottest month (see Appendix S3 for details).

Temporal variables were generated to represent the temporal stochastic processes in time (e.g., ecological drift) using the asymmetric eigenvector map (AEM) method, which is a filtering approach to model the effects produced by directed processes (e.g., time) (Blanchet et al. 2011). Studies of time-series dynamics often necessitate logical consistency, which posits that a previous point in time will affect a later point in time. Consequently, the AEM approach represents the optimal methodology for such studies (Baho et al. 2015). The sampling times of each sample point were sorted and placed in the same frame, and the AEMs were computed in such a way that each sampling time point corresponds to the corresponding AEM. In calculating the AEM, the sampling time interval was employed to determine the weights. The formula was as follows:

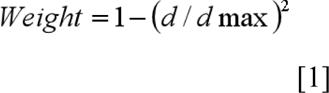

In this context, the notation “d” represents the sampling time interval, while “dmax” denotes the maximum value of the sampling time interval. Given a sample size of n, the AEM produces a total of n-1 positive eigenroots, with no negative eigenroots. The eigenfunctions can be divided into two sets of variables with positive and negative temporal correlation (Blanchet et al. 2008). Based on the Moran I coefficients, we selected positively correlated AEMs (Blanchet et al. 2011), as detailed in Appendix S3. The AEMs were computed with the “aem.time” function from the “adespatial” package (Dray 2022).

### 2.3 Data analysis

The objective of this study is to examine the spatial asymmetry of the environment and its variability. Principal component analysis (PCA) was conducted using the “FactoMineR” package in R (Lê et al. 2008) to examine the environmental variables’ averages and coefficients of variation (CV) over the study period. This was done in order to explore the principal axes of the environment and its variability. The objective of principal component analysis (PCA) is to reduce the dimensionality of a data set by transforming the data into a smaller number of principal components while retaining as much information as possible about the main sources of variation. This process not only contributes to the downscaling and simplification of the data, but also effectively reveals the underlying structure among the original variables and their interrelationships (Liu et al. 2024). The principal component coordinates were calculated and obtained through principal component analysis. Based on the eigenvalues, the first two principal component axes (PC1 and PC2) were selected, as they capture the majority of the variability in the data. Subsequently, the relationship between elevation and the two principal component axes was investigated using the generalized additive model (GAM) from the “mgcv” package in R (Wood et al. 2016).

The GAM model was employed to investigate elevation patterns in species richness across sample sites over 21 years. Subsequently, the presence and absence data were utilized to calculate three distance matrices for each sample point based on the Jaccard coefficient of variation. These matrices represent temporal beta diversity (Btotal), its decomposed species substitution (Brepl), and richness difference (Brich). The distance matrices were obtained using the “BAT” package in R, as outlined by Cardoso (2022). Subsequently, the GAM model was employed to investigate the elevation patterns of temporal beta diversity and its components, with the relative importance of the different components of temporal beta diversity then calculated. The Wilcoxon test was employed to ascertain which of the components of temporal beta diversity (Brep and Brich) was the primary driver.

To validate how elevational change in the relative significance of stochastic and deterministic processes can drive the shift of community dynamics, we employed generalized dissimilarity models (GDMs) to estimate temporal beta diversity and its components, while accounting for environmental and temporal variables (Ferrier et al. 2002, Ferrier et al. 2007). GDMs transform environmental variables through an iterative maximum likelihood fitting process to identify the optimal fit between environmental discrepancies and community differences (Jones et al. 2013).

The GDM assumes that biome variance increases monotonically with environmental variance and employs a link function to transform linear predictors in order to adjust for the nonlinear relationship between the two (Jones et al. 2013). The GDM model fit is constrained by the number of i-spline basis functions, with a default value of three i-splines (Fitzpatrick 2022), in instances where the model fit was not feasible, the number of i-splines was increased gradually until a maximum of five was reached. We conducted variance decomposition at each sample point to assess the rate of explanation of temporal beta diversity and its components by environmental and temporal variables. This was accomplished using the “gdm.partition.deviance” function in the R package “gdm” (Borcard et al. 1992, Jones et al. 2013). Then, the GAMs were employed to investigate the relationship between the ratio of pure explanatory rates of environmental and temporal variables and elevation, to identify patterns of elevation on the various ecological processes that shape community dynamics.

## 3 Results

### 3.1 Spatial asymmetry of the environment and its variability

The spatial distribution of averages and coefficients of variation of environmental variables exhibited a comparable pattern. The results of the principal component analysis demonstrated that Axis I and Axis II collectively explained 37.01% and 19.81% of the averages of the environmental variables, respectively (Fig. 2a). Additionally, 38.71% and 15.85% of the coefficient of variation of environmental variables were explained by axes I and II, respectively (Fig. 2b). For instance, the high-elevation sample point (1039 m) was markedly distant from other sample points, with a notable discrepancy.

**Figure 2.**
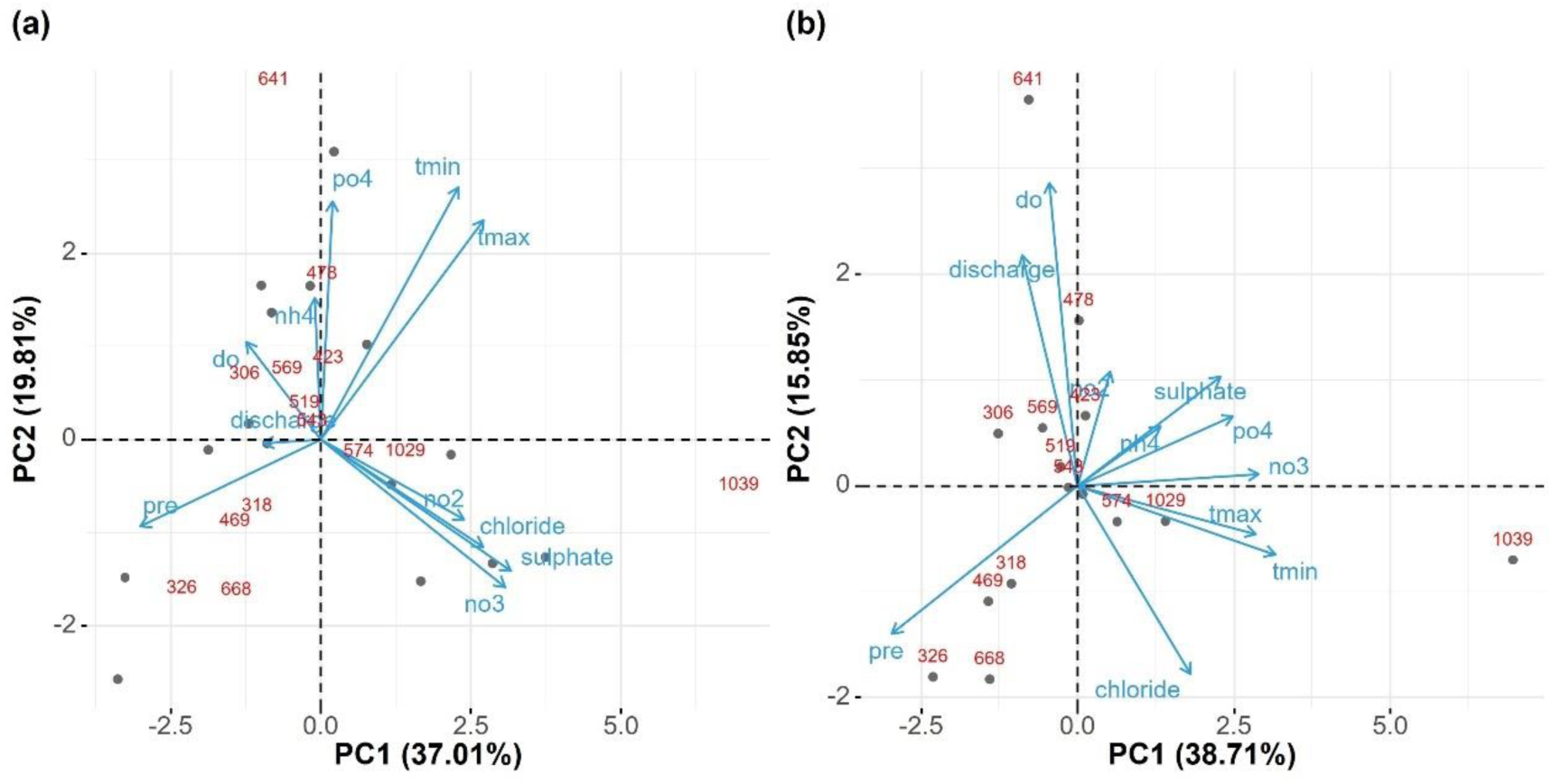
Biplots of principal component analysis (PCA) results for environmental variables, displaying (a) averages and (b) coefficients of variation (CV) over the study period.

In accordance with the PCA results pertaining to the averages of environmental variables, the environmental variables that exceeded the mean contribution to PC1, in descending order, were sulphates, nitrates, monthly mean precipitation, monthly mean maximum temperature, chlorides, ammonium, and monthly mean minimum temperature (for further details, please refer to Appendix 1, where the environmental variables that exceeded the average contribution were listed in descending order of their respective rankings). The environmental variables that contributed more to PC2 were monthly mean minimum temperature, phosphorus, and monthly mean maximum temperature. The PCA results for environmental variables demonstrated no significant differences between the two axes. Both axes were found to be related to climatic variables and water quality, but with axis one focusing on water quality and axis two on climatic variables.

The PCA results for the coefficient of variation of environmental variables indicated that the variables contributing the most to PC1 were monthly mean minimum temperature, monthly mean precipitation, nitrates, monthly mean maximum temperature, phosphorus, and sulphates. The variables that contributed more to PC2 were oxygen, discharge, and chlorides. The results of the principal component analysis (PCA) for the coefficient of variation of environmental variables indicated that axis one was associated with fluctuations in both water quality and climate variables, with climate fluctuations exerting a more pronounced influence. In contrast, axis two was closely linked to fluctuations in water quality.

The generalized additive model revealed a significant elevation pattern in the Axis I and Axis II scores of the averages of environmental variables, indicating a gradient change in the environment (water quality and climate) with elevation (Fig. 3a and 3c). While a significant elevation pattern was evident in the Axis I scores of the coefficient of variation of environmental variables (Fig. 3b), no such pattern was discernible in the Axis II scores (Fig. 3d). This indicated that the coefficient of variation of environmental variables demonstrated a certain degree of gradient with elevation. The evidence suggested that the coefficient of variation of climate exerted a greater influence than water quality. However, it should be noted that some the coefficient of variation of water quality do not result in a discernible change in gradient.

**Figure 3.**
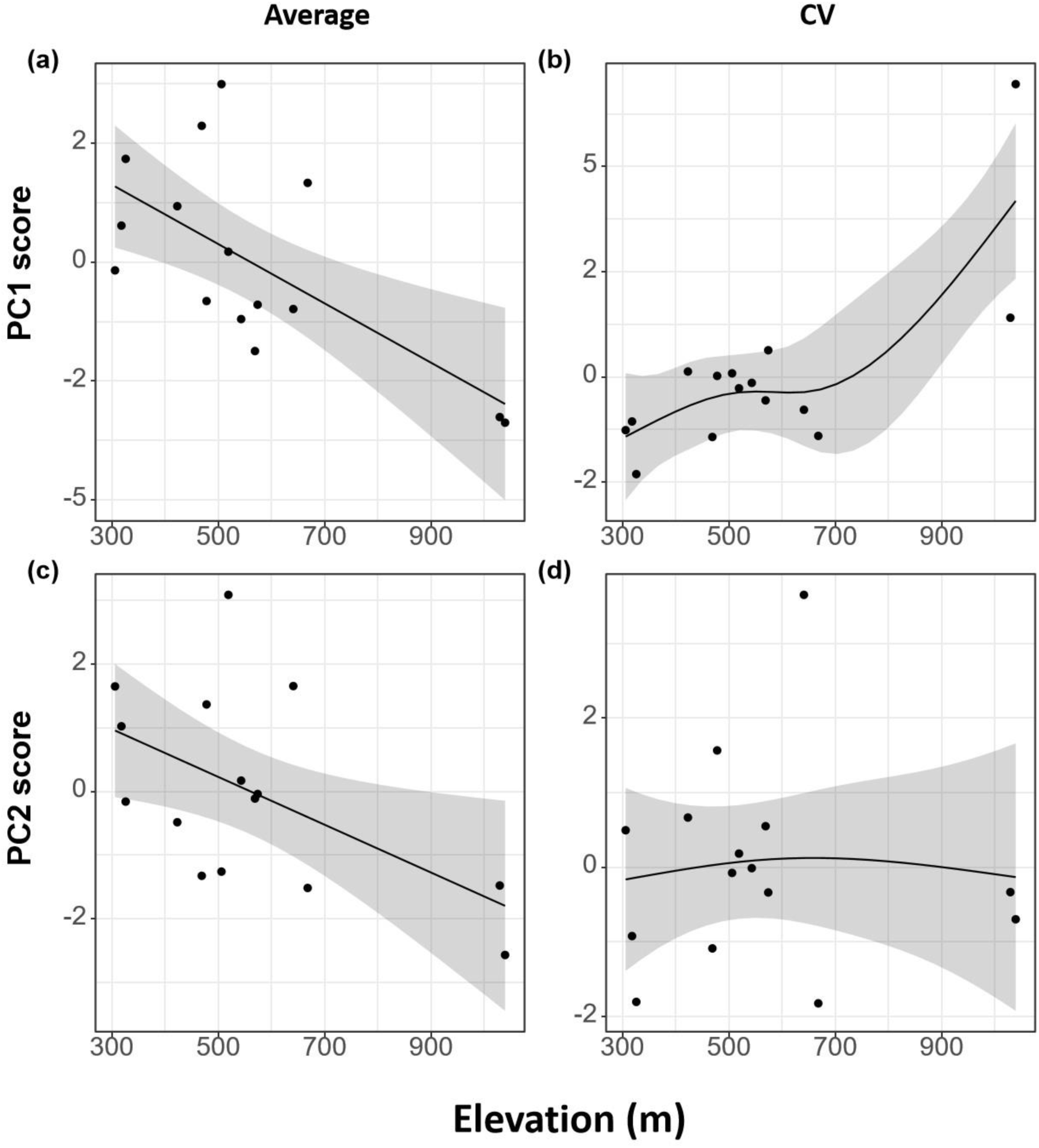
The elevation patterns of (a) axis one of the averages of environment variables, (b) axis one of the coefficients of variation (CV) of environment variables, (c) axis two of the averages of environment variables, (d) axis two of the coefficients of variation (CV) of environment variables. The patterns were analyzed using Generalized Additive Modeling.

### 3.2 Biotic dynamics

Using generalized additive modeling (GAM), species richness followed a bimodal pattern with elevation over the study period (Fig. 4a). The temporal beta diversity (Btoal) gradually decreased with increasing elevation, but its components Brepl and Brich did not show a clear or regular change with elevation (Fig. 4b, 4c and 4d). Species turnover (Brepl) was significantly higher than richness difference (Brich) when a Wilcoxon test for Brepl and Brich was performed. This result suggested that species turnover dominated the temporal beta diversity in the region (Refer to Appendix S1 for further details).

**Figure 4.**
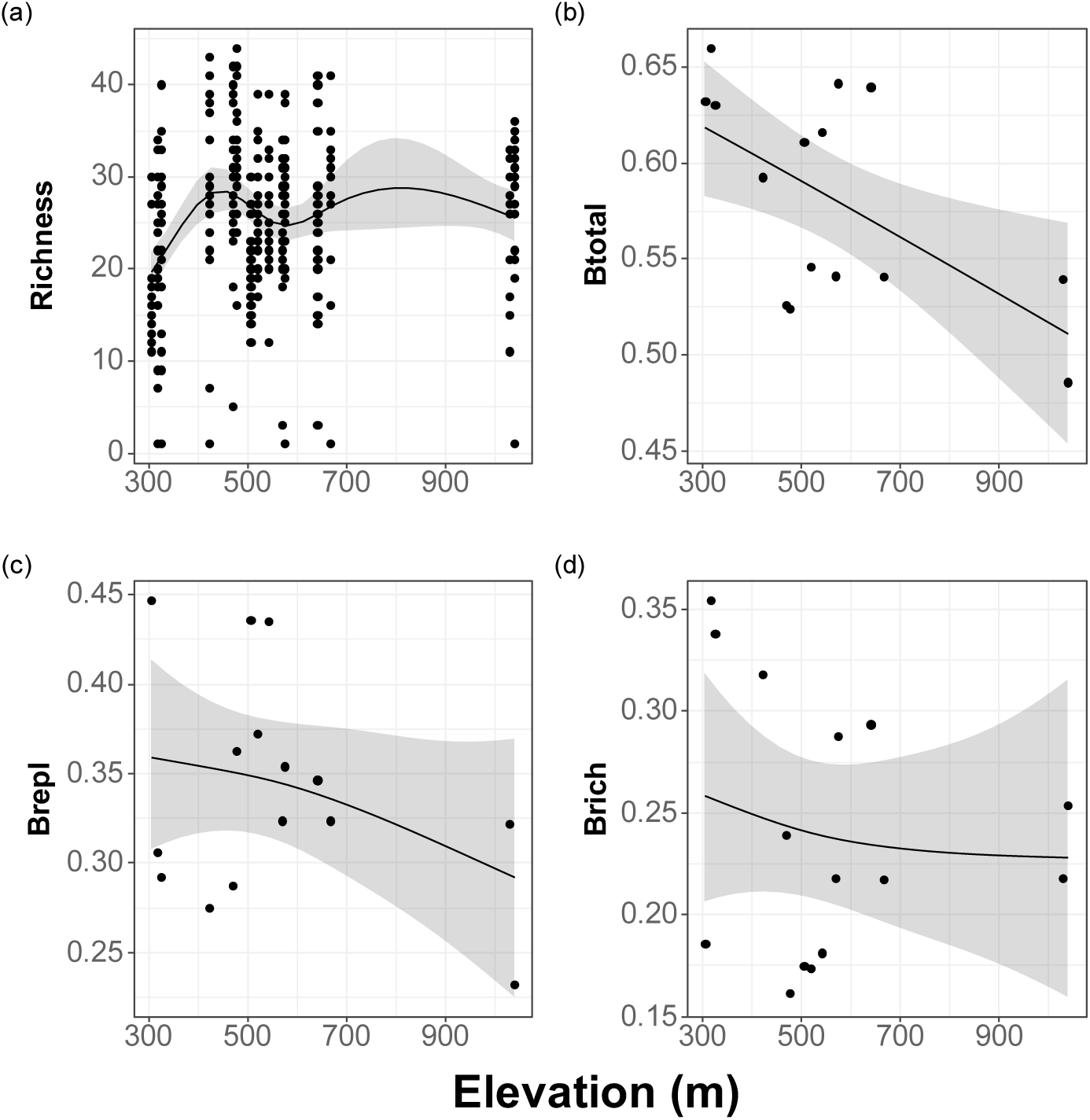
Elevation patterns of (a) Richness, (b) temporal beta diversity (Btotal), (c) species replacement (Brepl), and (d) richness difference in temporal beta diversity (Brich).

### 3.3 Ecological processes

The results demonstrated that no discernible pattern of elevation was evident for either Btotal or Brepl, regardless of whether deterministic or stochastic processes were considered. In the case of Brich, while no significant pattern of elevation was observed for either the deterministic or stochastic processes (Fig. 5f and 5i), a significant pattern of elevation was identified in the relative importance of these processes. As elevation increased, the relative importance of the processes in question appeared to diminish (Fig. 5c).

**Figure 5.**
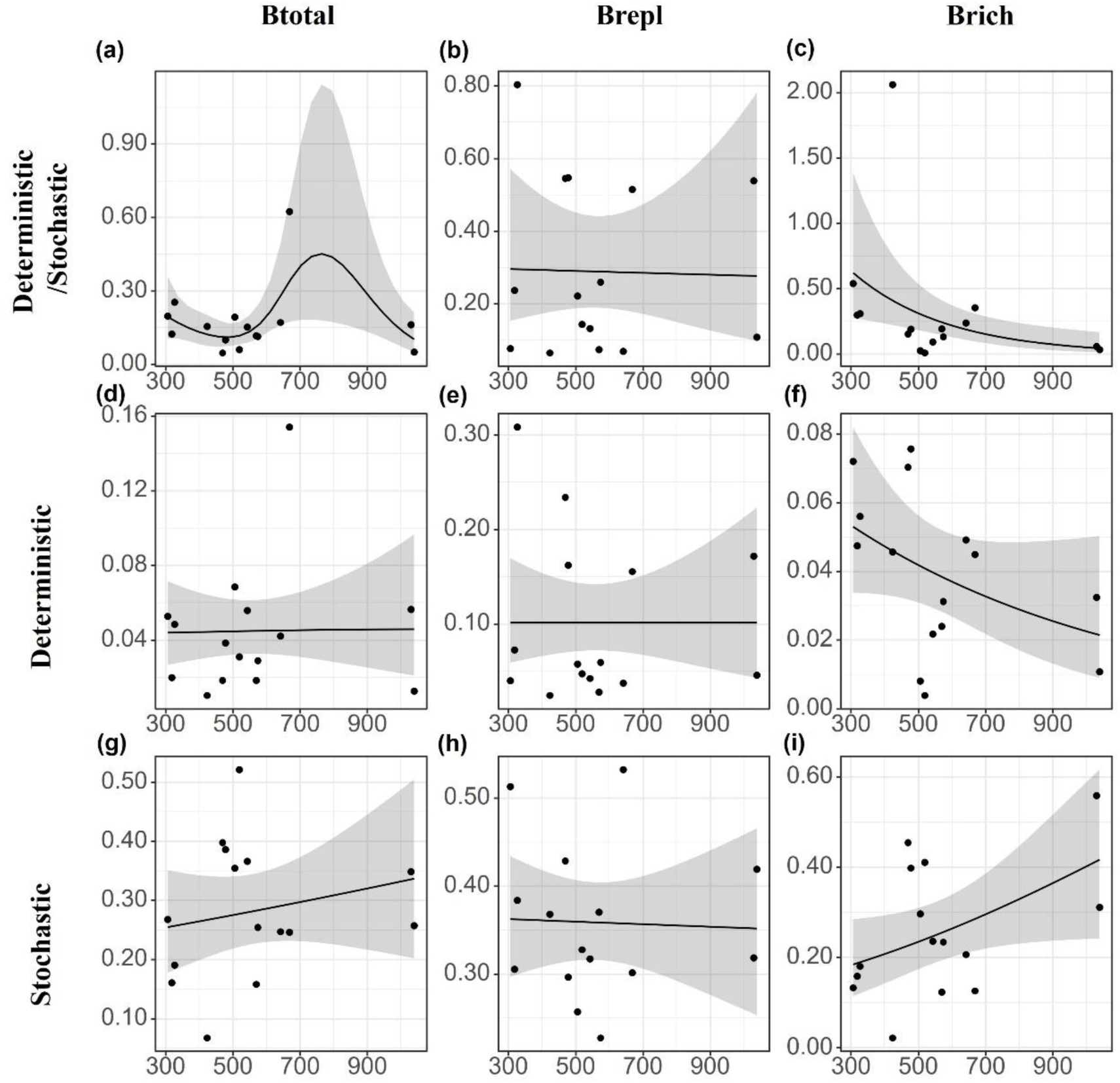
Elevation patterns of the relative importance of different ecological processes driving (a) temporal beta diversity (Btotal), (b) species replacement in temporal beta diversity (Brepl), and (c) richness difference in temporal beta diversity (Brich). Elevation patterns of the relative importance of climate driving (d) Btotal, (e) Brepl, and (f) Brich. Elevation patterns of the relative importance of AEM driving (g) Btotal, (h) Brepl, and (i) Brich.

## 4 Discussion

An investigation was conducted to examine the geographic patterns in community dynamics and environments variability using 21 years of long-term data on benthic macroinvertebrates. To gain further insight into the underlying ecological mechanisms, we linked these patterns to temporal ecological processes and discovered that the geographic asymmetry in environmental variability shaped the spatial patterns of community dynamics and temporal ecological processes. The hypothesis that the importance of ecological processes exhibited a gradient with space was supported by the findings. The relative importance of deterministic versus stochastic processes decreased with increasing elevation.

### 4.1 Spatial variation of community dynamics

Our findings revealed a partial bimodal pattern of macroinvertebrates richness, which is constrained by the elevational gradient of the study area. Elevational patterns of species richness demonstrated a diverse pattern as the study taxa and study area changed (Harrington et al. 2016, Wei et al. 2024). Temporal beta diversity gradually declined with increasing elevation, suggesting that macroinvertebrate community fluctuations gradually decreased and community stability increased. A growing body of evidence suggested that as species richness increased, the community became more stable, which is an example of the “portfolio effect”, where the stability of the community increases with diversity. This finding has been extensively supported by studies of the diversity-stability ratio (DSR) of ecosystems (Campbell et al. 2011). For instance, with increasing macroinvertebrates richness, the temporal synchronization of populations in a community diminishes, indicating that the community became more resistant to fluctuations and became more stable (Roscher et al. 2011). Nevertheless, our findings are not aligned with those of the aforementioned study. The discrepancy in the results may be attributed to the influence of macroinvertebrates abundance. Communities with greater abundance generally have fewer fluctuations in the community, and the distribution of population-abundance heterogeneity also impacts DSR (Kilpatrick and Ives 2003, Thibaut and Connolly 2013).

The results of this study indicated that temporal beta diversity was primarily driven by species replacement, which is consistent with previous findings (Angeler 2013, Tonkin et al. 2017, Wu et al. 2022). The frequent turnover of macroinvertebrate species may be attributed to the high temporal heterogeneity of the environment at the study site. Environmental change plays a crucial role in the worldwide distribution of biodiversity (Jocque et al. 2010, Milner et al. 2023). The region’s pronounced environmental fluctuations have introduced novel temporal ecological niches for macroinvertebrates, thereby promoting local species survival and facilitating species turnover (Heino and Tolonen 2017, Tonkin et al. 2017, Wu et al. 2022). Additionally, macroinvertebrates possess unique life history responses and evolutionary strategies that allow them to exploit these temporal ecological niches, potentially leading to significant species turnover (Wu et al. 2022). For example, Baetidae were more adapted to high-flow stream environments and exhibited heightened activity during the spring and summer months, when water levels rose due to warmer temperatures and increased oxygenation. In contrast, species belonging to Ephemeridae were more frequently observed during the autumn and winter months, when conditions were cooler and water flow was reduced (Merritt et al. 2019). The rapid turnover observed among macroinvertebrates often coincided with changes in rare taxa (Mori et al. 2010, Rocha et al. 2019, Castro et al. 2020), which are particularly sensitive and vulnerable to environmental fluctuations (Monaghan et al. 2005, Nyirabuhoro et al. 2020). For example, Anthomyiidae were highly susceptible to variability in temperature and humidity, while Astacidae demonstrated a notable sensitivity to alterations in dissolved oxygen levels and the presence of pollutants, including heavy metals and organic matter (Merritt et al. 2019). The frequency of species occurrence during the study period is detailed in Appendix S1.

### 4.2 Inconsistent ecological processes across the landscape

Our findings indicated that stochastic and deterministic processes, when considered together, exerted an influence on the dynamics of macroinvertebrate communities in the study area. For Brich, the relative importance of deterministic versus stochastic processes was found to decrease with increasing altitude, with stochastic processes ultimately exerting a dominant influence. These findings were likely associated with resource scarcity, which resulted in an environmental gradient as elevation increased. In environments with limited resources, the influence of deterministic processes (such as macroinvertebrate interactions) was diminished. This was evident in competitive interactions between Asellidae and Gammaridae, as well as predation by Dytiscidae on larvae belonging to Chironomidae. This allowed stochastic events in ecosystems, such as climate change, habitat disturbance, or sudden changes in food availability, to have a significant influence on survival and reproduction. Consequently, there was an increased likelihood of stochastic factors influencing community dynamics (Conradi et al. 2017, Yang et al. 2024).

It was suggested that these results were closely related to environmental variability, which exhibited a gradient across elevations. When environmental variability fell within the tolerance range of macroinvertebrates, macroinvertebrates were filtered more strongly as the variability increased, having a greater impact on communities (Vorste et al. 2016). The capacity to respond to environmental variation differed between different ecological roles. For instance, Baetidae, which were herbivores, exhibited a greater sensitivity to temperature than Perlidae, which were predators (Shah et al. 2021). However, once a certain threshold was exceeded, heightened variability could lead to macroinvertebrates mortality, resetting the community and introducing stochastic processes such as stochastic colonization, further emphasizing the importance of these processes (Chase 2003). Frequent extreme climatic events led to stochastic patterns of species extinction and colonization, and diminished the resilience of ecosystems. The frequency, intensity, and periodicity of these events, including floods and droughts, had a significant impact on community dynamics (Chiu et al. 2021). Such disturbances had the potential to disrupt the return of macroinvertebrate communities to their optimal ecological niche, thereby reducing interspecific competition. Therefore, these disturbances manifested as an amplification of stochastic processes, which obscured spatial and temporal patterns.

### Conclusions

Our findings underscore the necessity of differentiating between temporal community dynamics and observed spatial variation to optimize local biodiversity management and conservation initiatives across diverse landscapes. The influence of spatial asymmetries in the environment, including water quality and climate fluctuations, on the changes in ecological mechanisms driving community dynamics is a significant factor that must be considered. The analysis of spatial heterogeneity in temporal ecological processes can provide valuable insights into the distribution of biodiversity. This can inform predictions regarding the geographical migration of organisms in order to adapt to an environment of increased variability under global warming scenarios. It is our contention that further research will provide a robust theoretical foundation for predicting the geographic distribution of community dynamics. It is therefore imperative that long-term biomonitoring be maintained, particularly in regions exhibiting pronounced environmental variability, such as mountainous areas. In response to spatial asymmetries in community dynamics under environmental variability, the predictability and effectiveness of regional conservation strategies can be significantly enhanced by the implementation of locally appropriate biodiversity conservation measures.

## Acknowledgments

We gratefully acknowledge the authors (Canedo-Arguelles, M., Gutierrez-Canovas, C., Acosta, R., Castro-Lopez, D., Cid, N., Fortuno, P., et al.) of the study titled “As time goes by: 20 years of changes in the aquatic macroinvertebrate metacommunity of Mediterranean river networks” for generously sharing their data.

## References

Aguilar, P., and R. Sommaruga. 2020. The balance between deterministic and stochastic processes in structuring lake bacterioplankton community over time. Molecular Ecology 2: 3117–3130.

Angeler, D. G. 2013. Revealing a conservation challenge through partitioned long-term beta diversity: increasing turnover and decreasing nestedness of boreal lake metacommunities. Diversity and Distributions 1: 772–781.

Antunez, P. 2023. Evidence of the variation in the rate of change of temperature and precipitation. Ecological Informatics.

Baho, D. L., M. N. Futter, R. K. Johnson, and D. G. Angeler. 2015. Assessing temporal scales and patterns in time series: Comparing methods based on redundancy analysis. Ecological Complexity 22:162–168.

Bier, R. L., M. Vass, A. J. Székely, and S. Langenheder. 2022. Ecosystem size-induced environmental fluctuations affect the temporal dynamics of community assembly mechanisms. The ISME Journal 1: 2635–2643.

Blanchet, F. G., P. Legendre, and D. Borcard. 2008. Modelling directional spatial processes in ecological data. Ecological Modelling 21: 325–336.

Blanchet, F. G., P. Legendre, R. Maranger, D. Monti, and P. Pepin. 2011. Modelling the effect of directional spatial ecological processes at different scales. Oecologia 1: 357–368.

Bontemps, Z., Y. Moenne-Loccoz, and M. Hugoni. 2024. Stochastic and deterministic assembly processes of microbial communities in relation to natural attenuation of black stains in Lascaux Cave. Msystems.

Borcard, D., P. Legendre, and P. Drapeau. 1992. Partialling out the spatial component of ecological variation. Ecology: 1045–1055.

Campbell, V., G. Murphy, and T. N. Romanuk. 2011. Experimental design and the outcome and interpretation of diversity-stability relations. Oikos 12: 399–408.

Cañedo-Argüelles, M., C. Gutiérrez-Cánovas, R. Acosta, D. Castro-López, N. Cid, P. Fortuño, A. Munné, C. Múrria, A. R. Pimentao, R. Sarremejane, M. Soria, P. Tarrats, I. Verkaik, N. Prat, and N. Bonada. 2020. As time goes by: 20 years of changes in the aquatic macroinvertebrate metacommunity of Mediterranean river networks. Journal of Biogeography: 1861–1874.

Cardoso, P. M., S.; Rigal, F.; Carvalho, J. 2022. BAT: Biodiversity Assessment Tools.

Castro, D. M. P., P. G. da Silva, R. Solar, and M. Callisto. 2020. Unveiling patterns of taxonomic and functional diversities of stream insects across four spatial scales in the neotropical savanna. Ecological Indicators 11: 8.

Chan, W. P., I. C. Chen, R. K. Colwell, W. C. Liu, C. Huang, and S. F. Shen. 2018. Seasonal and daily climate variation have opposite effects on species elevational range size (vol 351, pg 1437, 2016). Science.

Chase, J. M. 2003. Community assembly: when should history matter? Oecologia 1: 489–498.

Chiu, M. C., S. C. Ao, V. H. Resh, F. Z. He, and Q. H. Cai. 2021. Species dispersal along rivers and streams may have variable importance to metapopulation structure. Science of the Total Environment.

Conradi, T., V. M. Temperton, and J. Kollmann. 2017. Resource availability determines the importance of niche-based versus stochastic community assembly in grasslands. Oikos 12: 1134–1141.

Deng, Y. 2023. mixgb: Multiple Imputation Through ‘XGBoost’.

Dini-Andreote, F., J. C. Stegen, J. D. van Elsas, and J. F. Salles. 2015. Disentangling mechanisms that mediate the balance between stochastic and deterministic processes in microbial succession. Proceedings of the National Academy of Sciences of the United States of America 112:E1326–E1332.

Doxa, A., Y. Kamarianakis, and A. D. Mazaris. 2022. Spatial heterogeneity and temporal stability characterize future climatic refugia in Mediterranean Europe. Global Change Biology 2: 2413–2424.

Dray, S. B., D.; Blanchet, G.; Borcard, D.; Clappe, S.; Guénard, G.; Jombart, T.; Larocque, G.; Legendre, P.; Madi, N.; Wagner, H.H. 2022. adespatial: Multivariate Multiscale Spatial Analysis.

Ferrier, S., M. Drielsma, G. Manion, and G. Watson. 2002. Extended statistical approaches to modelling spatial pattern in biodiversity in northeast New South Wales. II. Community-level modelling. Biodiversity and Conservation 11:2309–2338.

Ferrier, S., G. Manion, J. Elith, and K. Richardson. 2007. Using generalized dissimilarity modelling to analyse and predict patterns of beta diversity in regional biodiversity assessment. Diversity and Distributions 1: 252–264.

Fick, S. E., and R. J. Hijmans. 2017. WorldClim 2: new 1-km spatial resolution climate surfaces for global land areas. International Journal of Climatology: 4302–4315.

Fitzpatrick, M. M., K.; Manion, G.; Nieto-Lugilde, D.; Ferrier, S. 2022. gdm: Generalized Dissimilarity Modeling.

Gao, J., and Y. H. Liu. 2018. Climate stability is more important than water-energy variables in shaping the elevational variation in species richness. Ecology and Evolution: 6872–6879.

Harrington, R. A., N. L. Poff, and B. C. Kondratieff. 2016. Aquatic insect β-diversity is not dependent on elevation in Southern Rocky Mountain streams. Freshwater Biology 1:195–205.

He, X. L., C. Brown, and L. X. Lin. 2021. Relative importance of deterministic and stochastic processes for beta diversity of bird assemblages in Yunnan, China. Ecosphere 12.

Heino, J., and K. T. Tolonen. 2017. Ecological drivers of multiple facets of beta diversity in a lentic macroinvertebrate metacommunity. Limnology and Oceanography 2:2431–2444.

Hijmans, R. J. P., S.; Leathwick, J.; Elith, J. 2023. dismo: Species Distribution Modeling.

Hiltunen, T., G. B. Ayan, and L. Becks. 2015. Environmental fluctuations restrict eco-evolutionary dynamics in predator–prey system. Proceedings of the Royal Society B: Biological Sciences 2 2:20150013.

Jocque, M., R. Field, L. Brendonck, and L. De Meester. 2010. Climatic control of dispersal-ecological specialization trade-offs: a metacommunity process at the heart of the latitudinal diversity gradient? Global Ecology and Biogeography 1: 244–252.

Jones, M. M., S. Ferrier, R. Condit, G. Manion, S. Aguilar, and R. Perez. 2013. Strong congruence in tree and fern community turnover in response to soils and climate in central Panama. Journal of Ecology 1 1:506–516.

Kilpatrick, A. M., and A. R. Ives. 2003. Species interactions can explain Taylor’s power law for ecological time series. Nature 22:65–68.

Kozlovsky, D. Y., C. L. Branch, A. M. Pitera, and V. V. Pravosudov. 2018. Fluctuations in annual climatic extremes are associated with reproductive variation in resident mountain chickadees. Royal Society Open Science.

Lê, S., J. Josse, and F. Husson. 2008. FactoMineR: An R Package for Multivariate Analysis. Journal of Statistical Software 2: 1–18.

Lin, X., Z. Tian, Q. Luo, J. Li, Q. Cai, M.-C. Chiu, and V. H. Resh. 2024a. Spatial asymmetry of temporal ecological processes can shift in riverine macroinvertebrates responding to fluctuating climate conditions. Science of the Total Environment 2:175872.

Lin, X., J. Wan, Q. Luo, Q. Cai, M.-C. Chiu, and V. H. Resh. 2024b. Spatial Heterogeneity of Climate Fluctuations Shapes Ecological Processes and Community Dynamics in Riverine Invertebrates Across a Landscape. bioRxiv:2024.2008.2030.610482.

Liu, D., A. Esquivel-Muelbert, N. Acil, J. Astigarraga, E. Cienciala, J. Fridman, G. Kunstler, T. J. Matthews, P. Ruiz-Benito, J. P. Sadler, M.-J. Schelhaas, S. Suvanto, A. Talarczyk, C. W. Woodall, M. A. Zavala, C. Zhang, and T. A. M. Pugh. 2024. Mapping multi-dimensional variability in water stress strategies across temperate forests. Nature Communications 1: 8909.

Liu, X. J., N. Ferreira-Rodríguez, R. W. Wu, S. Ouyang, and X. P. Wu. 2023. Sixty years of species diversity and population density decline of freshwater mussels in a global biodiversity hotspot. Global Ecology and Conservation.

Magurran, A. E., M. Dornelas, F. Moyes, and P. A. Henderson. 2019. Temporal β diversity-A macroecological perspective. Global Ecology and Biogeography 2: 1949–1960.

Merritt, R. W., K. W. Cummins, and M. B. Berg. 2019. An introduction to the aquatic insects of North America (Fifth edition). Kendall Hunt Publishing Company.

Milner, A. M., E. M. L. Vega, T. J. Matthews, S. C. Conn, and F. M. Windsor. 2023. Long-term changes in macroinvertebrate communities across high-latitude streams. Global Change Biology 2: 2466–2477.

Monaghan, M. T., C. T. Robinson, P. Spaak, and J. V. Ward. 2005. Macroinvertebrate diversity in fragmented Alpine streams: implications for freshwater conservation. Aquatic Sciences: 454–464.

Mori, T., M. Murakami, and T. Saitoh. 2010. Latitudinal gradients in stream invertebrate assemblages at a regional scale on Hokkaido Island, Japan. Freshwater Biology: 1520–1532.

Nyirabuhoro, P., M. Liu, P. Xiao, L. M. Liu, Z. Yu, L. N. Wang, and J. Yang. 2020. Seasonal Variability of Conditionally Rare Taxa in the Water Column Bacterioplankton Community of Subtropical Reservoirs in China. Microbial Ecology: 14–26.

Rocha, M. P., L. M. Bini, M. Gronroos, J. Hjort, M. Lindholm, S.-M. Karjalainen, K. E. Tolonen, and J. Heino. 2019. Correlates of different facets and components of beta diversity in stream organisms. Oecologia 1 1:919–929.

Roscher, C., A. Weigelt, R. Proulx, E. Marquard, J. Schumacher, W. W. Weisser, and B. Schmid. 2011. Identifying population- and community-level mechanisms of diversity-stability relationships in experimental grasslands. Journal of Ecology: 1460–1469.

Sanchez-Montoya, M. M., M. R. Vidal-Abarca, T. Punti, J. M. Poquet, N. Prat, M. Rieradevall, J. Alba-Tercedor, C. Zamora-Munoz, M. Toro, S. Robles, M. Alvarez, and M. L. Suarez. 2009. Defining criteria to select reference sites in Mediterranean streams. Hydrobiologia 1: 39–54.

Shah, A. A., H. A. Woods, J. C. Havird, A. C. Encalada, A. S. Flecker, W. C. Funk, J. M. Guayasamin, B. C. Kondratieff, N. L. Poff, S. A. Thomas, K. R. Zamudio, and C. K. Ghalambor. 2021. Temperature dependence of metabolic rate in tropical and temperate aquatic insects: Support for the Climate Variability Hypothesis in mayflies but not stoneflies. Global Change Biology 2: 297–311.

Thibaut, L. M., and S. R. Connolly. 2013. Understanding diversity-stability relationships: towards a unified model of portfolio effects. Ecology Letters 1: 140–150.

Tonkin, J. D., M. T. Bogan, N. Bonada, B. Rios-Touma, and D. A. Lytle. 2017. Seasonality and predictability shape temporal species diversity. Ecology: 1201–1216.

Vanwonterghem, I., P. D. Jensen, P. G. Dennis, P. Hugenholtz, K. Rabaey, and G. W. Tyson. 2014. Deterministic processes guide long-term synchronised population dynamics in replicate anaerobic digesters. Isme Journal: 2015–2028.

Vorste, R. V., R. Corti, A. Sagouis, and T. Datry. 2016. Invertebrate communities in gravel-bed, braided rivers are highly resilient to flow intermittence. Freshwater Science: 164–177.

Wayman, J. P., J. P. Sadler, T. A. M. Pugh, T. E. Martin, J. A. Tobias, and T. J. Matthews. 2022. Assessing taxonomic and functional change in British breeding bird assemblages over time. Global Ecology and Biogeography 1:925–939.

Wei, M., T. Feng, Y. Lin, S. He, H. Yan, R. Qiao, and Q. Chen. 2024. Elevation-associated pathways mediate aquatic biodiversity at multi-trophic levels along a plateau inland river. Water Research 2: 121779.

Whitenack, L. E., J. F. Welklin, C. L. Branch, B. R. Sonnenberg, A. M. Pitera, D. Y. Kozlovsky, L. M. Benedict, V. K. Heinen, and V. V. Pravosudov. 2023. Complex relationships between climate and reproduction in a resident montane bird. Royal Society Open Science 1.

Wood, S. N., N. Pya, and B. Säfken. 2016. Smoothing Parameter and Model Selection for General Smooth Models. Journal of the American Statistical Association 111:1548–1563.

Wu, N. C., Y. C. Wang, Y. X. Wang, X. M. Sun, C. Faber, and N. Fohrer. 2022. Environment regimes play an important role in structuring trait-and taxonomy-based temporal beta diversity of riverine diatoms. Journal of Ecology 11: 1442–1454.

Yang, Z., J. Xu, J. Li, L. He, H. Xu, X. Guo, S. Xue, and Y. Cao. 2024. Stochastic Processes Shape Bacterial Community Diversity Patterns along Plant Niche Gradients. Agronomy 1: 204.

Zhang, Z., Y. Hautier, T. J. Bao, J. Yang, H. Qing, Z. L. Liu, M. Wang, T. K. Li, M. Yan, and G. L. Zhang. 2022. Species richness and asynchrony maintain the stability of primary productivity against seasonal climatic variability. Frontiers in Plant Science 1.

