## Appendix S1 for "Riverine Invertebrates Exhibit Spatial Asymmetry in Temporal Ecological Processes Across Landscapes"

**Table S1.** The elevation patterns of (a) axis one of the environment variables, (b) axis one of the fluctuations of environment variables, (c) axis two of the environment variables, (d) axis two of the fluctuations of environment variables.

|  |  | **Results** |
| --- | --- | --- |
| **PC1 of env** | **Intercept** | 1 |
|  | **p** | 0.007 ** |
|  | **Explained** | 43.60% |
| **PC2 of env** | **Intercept** | 1 |
|  | **p** | 0.036* |
|  | **Explained** | 29.60% |
| **PC1 of cv of env** | **Intercept** | 1 |
|  | **p** | 0.003** |
|  | **Explained** | 68.50% |
| **PC2 of cv of env** | **Intercept** | 1 |
|  | **p** | 0.870 |
|  | **Explained** | 4.73% |

**Table S2.** GAM models of elevation patterns of (a) Richness, (b) temporal beta diversity (Btotal), (c) species replacement (Brepl) and (d)richness difference in temporal beta diversity (Brich).

|  |  | **Results** |
| --- | --- | --- |
| **Richness** | **Intercept** | < 2 x 10^^-16^ |
|  | **p** | < 2 x 10^^-16^ |
|  | **Explained** | 9.80% |
| **Btotal** | **Intercept** | 1.09 x 10^^-11^ |
|  | **p** | 0.008 |
|  | **Explained** | 34.90% |
| **Brepl** | **Intercept** | < 2 x 10^^-16^ |
|  | **p** | 0.291 |
|  | **Explained** | 15.70% |
| **Brich** | **Intercept** | < 2 x 10^^-16^ |
|  | **p** | 0.721 |
|  | **Explained** | 6.09% |

**Table S3.** GAM models of elevation patterns of the relative importance of different ecological processes driving (a) temporal beta diversity (Btotal), (b) species replacement in temporal beta diversity (Brepl), and (c) richness difference in temporal beta diversity (Brich). Elevation patterns of the relative importance of water quality driving (d) Btotal, (e) Brepl, and (f) Brich. Elevation patterns of the relative importance of AEM driving (g) Btotal, (h) Brepl, and (i) Brich.

|  | | **Results** |
| --- | --- | --- |
| **Determinism /** **Stochasticity**  **(Btotal)** | Intercept | 3.49 x 10^^-8^ |
|  | p | 0.627 |
|  | Explained | 1.88% |
| **Determinism /** **Stochasticity**  **(Brepl)** | Intercept | 1.45 x 10^^-6^ |
|  | p | 0.652 |
|  | Explained | 8.43% |
| **Determinism /** **Stochasticity**  **(Brich)** | Intercept | 1.48 x 10^^-5^ |
|  | p | 0.096 |
|  | Explained | 14.60% |
| **Determinism (Btotal)** | Intercept | < 2 x 10^^-16^ |
|  | p | 0.533 |
|  | Explained | 3.07% |
| **Determinism (Brepl)** | Intercept | < 2 x 10^^-16^ |
|  | p | 0.958 |
|  | Explained | 0.16% |
| **Determinism (Brich)** | Intercept | < 2 x 10^^-16^ |
|  | p | 0.234 |
|  | Explained | 21.20% |
| **Stochasticity (Btotal)** | Intercept | 0.21 |
|  | p | 0.748 |
|  | Explained | 0.96% |
| **Stochasticity (Brepl)** | Intercept | 8.55 x 10^^-5^ |
|  | p | 0.981 |
|  | Explained | 0.05% |
| **Stochasticity (Brich)** | Intercept | 0.445 |
|  | p | 0.406 |
|  | Explained | 6.41% |

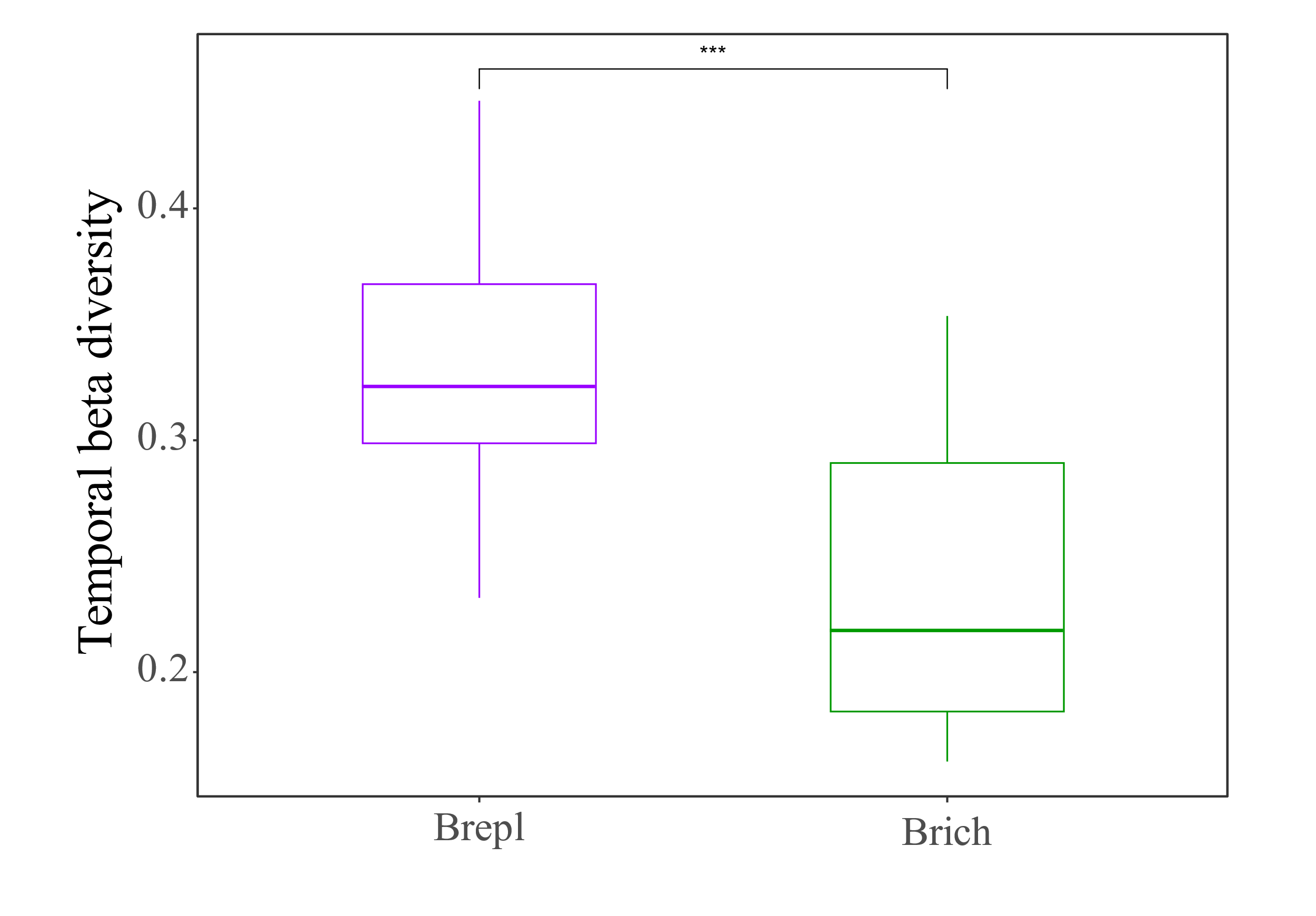

**Figure S1.** **The components of temporal beta diversity during the study period.**
